# Sialyl-Tn-positive tumour-derived extracellular vesicles impair dendritic cell function via horizontal transfer of glycans

**DOI:** 10.64898/2026.07.24.740563

**Authors:** Nayara Delgado André, Shally Sharma, Mariana Sousa Vieira, Brenda Raissa de Oliveira, Zélia Silva, Paula A Videira

## Abstract

The sialyl-Tn (STn) glycan antigen is aberrantly expressed in a subset of triple-negative breast cancer (TNBC) and is associated with poor prognosis and immunosuppressive microenvironment. Tumour-derived extracellular vesicles (TDEVs) are emerging regulators of immune escape however the role of glycan-mediated mechanisms remains elusive. Aberrant glycosylation is a hallmark of cancer that extends to TDEVs, yet how tumour-associated glycans within EV cargo modulate cell function remains poorly understood. Here we used engineered MDA-MB-231 TNBC cells to overexpress the glycosyltransferase ST6GalNAc-I, generating STn-positive cells whose EVs were enriched in STn (STn^+^ EV). The STn^+^ EVs impaired the maturation of monocyte-derived dendritic cells (DCs), reduced antigen presentation, and diminished CD4□ and CD8□ T cell priming, alongside the expansion of regulatory T cells. DCs co-cultured with STn□ EVs display STn at their cell surface. Notably, STn^+^ EVs transferred both STn antigen and the ST6GalNAc-I to recipient DCs. Enzymatic removal of terminal sialic acids from STn□ EVs reversed the immunosuppressive effects, confirming the STn□dependent nature of DC dysfunction.

These findings add STn to the extensive list of components of EVs’ molecular cargo that play a role in immune suppression and may contribute for developing precision medicine approaches in oncology.

## 1 Introduction

Cancer is a heterogeneous group of diseases characterized by dysregulated cell proliferation, impaired apoptosis, and the ability to invade and metastasize to distant organs, resulting from disruption in normal cellular growth and repair mechanisms (1,2). Despite significant advancements in surgery, chemotherapy, and immunotherapy, cancer remains a leading global cause of mortality. Breast cancer is the second most common malignancy among women, with more than 2.3 million new cases and approximately 670000 deaths reported in 2022 (3) High rate of breast cancer mortality is associated with late-stage diagnoses and limited treatments against metastasis (4–6). While genetic and molecular variations across different tumour cell types have already been extensively analysed, emerging evidence highlights the critical role of the tumour microenvironment (TME) in driving disease progression and resistance (7–9). TME is a dynamic and complex ecosystem comprising diverse components, including stromal elements, the extracellular matrix, signalling molecules, and immune cells (10,11). Among all these important components, the immune system plays a pivotal role, exerting multifaceted functions that promote and suppress tumour development in different organs (12,13).

Dendritic cells (DCs) are professional antigen-presenting cells (APCs) and play a critical role in initiating and regulating the adaptive immune response, making them central to the body’s defence against cancer (14–16). They stand out as key mediators of anti-tumour immunity, functioning as sentinels that constantly survey the body for abnormal cells (17–19). Paradoxically, tumours often exploit immune evasion mechanisms to facilitate their growth and dissemination (20,21), and in breast cancer patients, the function of dendritic cells is often compromised, contributing to immune evasion and tumour progression (22,23). In breast cancer, this immune-tumour interplay is particularly complex, as the balance between pro- and anti-tumourigenic immune responses can shift throughout the course of the disease (24,25).

In recent years, tumour-derived EVs (TDEVs) have emerged as critical mediators of intercellular communication within the TME (26–28). These nanosized (30 to 100 nm) EVs are secreted by both normal cells and prominently by tumour cells and carry active molecules, including proteins, lipids, nucleic acids, and glycans, which are involved in several physiological and pathological processes. EV’s cargo reflects the physiological state of the originating cell and can influence other cells by modulating their behaviour and function through intercellular communication mechanisms (29,30). TDEVs also influence immune surveillance by contributing to the presentation of tumour antigens and the activation of cytotoxic T cells. (31,32). Aberrant glycosylation is a common feature of cancer, and the altered cell-surface carbohydrates significantly influence tumour-immune interactions, modulating cell-cell communication, migration, and immune recognition, and contributing to tumour progression, angiogenesis, and immune evasion. (33–35). Tumour-associated glycans, such as Sialyl Tn, are particularly noteworthy as they are rarely expressed in normal tissues but are overexpressed in various malignancies, including breast cancer (36–38). STn is associated with elevated capacity for proliferation and metastasis, somatic cell reprogramming and stemness, epithelial-mesenchymal transition, and resistance to apoptosis and chemotherapy (36,39). Moreover, STn has been shown to have an immunosuppressive role in DC function (40). A recent study investigating STn’s role in TNBC (Triple negative breast cancer) detected STn in primary tumour tissues in 23.8% out of 126 TNBC patients, who exhibited significantly reduced survival and lower c-Myc expression. Additionally, data from The Cancer Genome Atlas (TCGA) TNBC confirmed this association, showing that high levels of *ST6GALNAC1,* the gene encoding the enzyme responsible for the STn synthesis, were inversely correlated with *MYC* expression and positively associated with TGF-β signalling genes and immunosuppressive cell infiltrates, such as macrophages M2 and regulatory T cells (41). Glycosylation has been implicated in protein sorting in EV formation (42). Moreover, an enhanced EV production was reported in STn antigen–expressing cells and the consequent augmented delivery of EVs to recipient cells (43). Despite the growing body of evidence on the importance TDEVs in tumour-immune cell crosstalk, the precise mechanisms underlying this interaction, as well as the role of EVs enriched with aberrant glycans, remain poorly understood. The present study aims to investigate the role of EVs bearing STn antigen in mediating intercellular communication between tumour cells and DCs, with an emphasis on their immunomodulatory effects. The impairment of DCs’ function in breast cancer patients has significant implications for both disease progression and therapeutic response (44,45). Effective immunotherapies, such as immune checkpoint inhibitors or cancer vaccines, rely on a functional DC-T cell axis to mount a robust antitumour response. Unfortunately, the dysfunction of DCs in the breast cancer microenvironment limits the success of these therapies in many patients (46–48). By elucidating whether and how EVs’ molecular cargo contributes to damping DC’s function sheds light on immune evasion mechanisms in breast cancer and may provide a foundation to restore the immune function and contribute to advancing novel immunotherapeutic approaches.

## 2 Materials and Methods

### 2.1 Cell Culture

Human breast cancer cell lines, MDA-MB-231 (ATCC® HTB-22™) wild-type (WT), MDA-MB-231 Mock transduced and MDA-MB-231 transduced to stably overexpress *ST6GALNAC1* to produce Sialyl Tn (MDA-MB-231-STn) (49) were used in this study. Cells were cultured in Dulbecco’s Modified Eagle Medium (DMEM) supplemented with 10% fetal bovine serum (FBS), 2 mM L-glutamine, and 1% penicillin-streptomycin (Thermo Fisher Scientific Gibco, USA). Cells were cultured at 37°C in a humidified incubator with 5% CO□ To evaluate EVs secretion in the supernatant, when cultures reached 60% cell confluency, the complete DMEM medium was replaced with DMEM containing ultracentrifuged FBS (uFBS), prepared by centrifuging FBS at 100 000× g for 4 h at 4°C. The supernatants were collected, once the cells reached 95% confluence.

### 2.2 Extracellular Vesicles Isolation

Extracellular vesicles were isolated from culture supernatants using a differential ultracentrifugation protocol with some modifications described by Abusamra (50). Briefly 50 ml of supernatant was centrifuged at 1 200×g for 10 min to remove cells and at 17 000× g for 45 min to remove cell debris. The resulting supernatant was ultracentrifuged at 100000× g for 110 min at 4°C in fixed-angle rotor (Beckman Coulter) and subsequently EV pellet was spin washed with phosphate-buffered saline (PBS). A final ultracentrifugation was done at 100 000× g for 110 minutes and the pellet was resuspended in 150 µL of PBS, filtered using 0.22 µm membrane (COSTAR® SPIN-X, Sigma) and stored at -80 °C for further analysis.

### 2.3 Transmission Electron Microscopy Analysis

Extracellular vesicles morphology and size were assessed by transmission electron microscopy (TEM) analysis. Samples were adsorbed onto 300-mesh carbon-coated copper grids (Lacey Formvar/Carbon; Electron Microscopy Sciences, USA) previously glow-discharged for 1 min. EVs were fixed with 1.5% glutaraldehyde for 10 min, followed by negative staining with 1% filtered uranyl acetate applied dropwise on the surface of the EM grid with the syringe. Excess uranyl acetate was removed using water and the grid was allowed to dry in dark for 20 min. The images were taken using TEM at a magnification of 49 000×

### 2.4 Nanoparticle Tracking Analysis

The size distribution and concentration of isolated EVs were determined using nanoparticle tracking analysis performed on the NanoSight LM10 system (Malvern Instruments, UK). The NTA instrumentation was configured with a laser of 488 nm with high sensitivity and resolution sCMOS camera. The EV samples were diluted in 1:1 with filtered PBS and analysed with a constant flow rate at room temperature. The particle frame was adjusted for 20-100 particles per frame, and data was analysed using NTA 3.2 Build 3.2.16 software with a bin size of 2 and detection threshold of 10.

### 2.5 Sialidase Treatment of Extracellular Vesicles

To enzymatically remove terminal sialic acid residues, EVs derived from MDA-MB-231 cells were treated with *Clostridium perfringens* neuraminidase (Sigma-Aldrich, USA). A total of 30 μg of EV protein was incubated with 5 U sialidase in 200 μL of sodium acetate buffer (pH 5.5). The enzyme concentration and incubation parameters were optimized in preliminary assays to achieve desialylation without compromising vesicle integrity. After incubation, the reaction was terminated by dilution with cold PBS, and EVs were washed by ultracentrifugation (100 000× g, 110 min, 4°C) to remove residual enzyme. The resulting pellet was resuspended in PBS and filtered through a sterile 0.22μm membrane. Treated EVs were then used in monocyte-derived dendritic cell (DC) differentiation assays. Control EVs (non-treated) were processed in parallel under identical conditions in the absence of sialidase.

### 2.6 Lysate Preparation

Cells were harvested using Trypsin-EDTA solution (0.25%, 1X; Sigma-Aldrich, T4049), incubated for 3–5 minutes at 37°C, and washed twice with ice-cold phosphate-buffered saline (PBS, pH 7.4). Whole-cell and EV lysates were obtained following a standardized protocol to preserve protein integrity using PierceTM IP Lysis Buffer (Thermo Fisher Scientific) containing cOmpleteTM, Mini, EDTA-free Protease Inhibitor Cocktail (Roche, Basel, Switzerland). Lysates were subjected to three repeated cycles of vigorous vortexing (30 sec) followed by incubation on ice (5 min) to ensure complete membrane disruption. The lysates were clarified by centrifugation at 13000× g for 15 min at 4°C, and the supernatants were collected for protein quantification.

Protein was quantified with the PierceTM BCA Protein Assay kit (Thermo Fisher Scientific) following the manufacturer’s instructions.

### 2.7 Western Blot

Cell or EV lysates were separated in 10% SDS-PAGE, using 30 µg per lane, and transferred to PVDF membranes (GE Healthcare Life Sciences, USA). For general protein detection, blocking was performed with 5% BSA in Tris-buffered saline with 0.05% Tween 20% (TBS-T). For STn glycan detection, membranes were blocked with a carbohydrate-free blocking solution for 1h at room temperature. Membranes were incubated overnight at 4°C with primary monoclonal antibodies: rabbit polyclonal anti-ST6GalNAc1 antibody (Invitrogen, PA5-31200; 1:1000 dilution); mouse anti-human STn (51), L2A5, 1:2000); mouse anti-CD9 (BD Bioscience, 555370,1:1000); mouse anti-CD81 (BD Bioscience, 555675, 1:1000), mouse anti-β-actin (Santa Cruz, sc-81178, 1:2000). After three 5-min washes with washing buffer solution, membranes were incubated with HRP-conjugated secondary antibodies: anti-rabbit IgG secondary antibody (Abcam, ab6721, 1:5000) or goat anti-mouse IgG secondary antibody (Abcam, ab205719, 1:2500) for 1h at room temperature. Detection was performed through enhanced chemiluminescence (ECL) using a 1:1 mixture of two substrate solutions with signal development captured after 10min of exposure using a digital imaging system.

### 2.8 Immunofluorescence Staining of Sialyl-Tn in MDA-MB-231 Cells

Immunofluorescence was performed to evaluate Sialyl-Tn (STn) antigen expression in MDA-MB-231 Mock and STn-transduced cells. Cells were seeded onto glass coverslips, fixed with 4% paraformaldehyde for 15 min at room temperature, and permeabilized using 0.1% Triton X-100 in PBS for 10 min. Non-specific binding was blocked with 1% BSA in PBS for 1 h at room temperature. STn detection was carried out using mouse anti-human monoclonal antibody (51), L2A5, 1:50) as primary antibody and incubation overnight at 4°C. After washing with PBS, cells were incubated with goat anti-mouse IgG conjugated to Alexa Fluor fluorophores 488 (Thermo Fisher, A-11001, 1:100) for 1 h at room temperature in the dark. Nuclei were counterstained with DAPI (1µg/ml) for 5 min. Coverslips were mounted with antifade reagent and visualized using a fluorescence microscope (ZEISS Axio Imager 2, Germany). Image acquisition and analysis were performed under identical exposure conditions across experimental groups.

### 2.9 Quantitative Real Time PCR

To assess the expression of the sialyltransferases involved in the synthesis of STn, RNA was extracted from cells and EVs using the GenElute Mammalian Total RNA Miniprep Kit (Sigma-Aldrich, RTN70). The integrity and purity of RNA were assessed by spectrophotometric analysis using a Nanodrop device, considering A260/280 and A260/230 absorbance ratios. Ratios close to 2.0 and 2.0–2.2 respectively, indicate high purity and absence of significant protein or solvent contamination in the extracted samples and 1 µg of total RNA was reverse-transcribed into cDNA using the High-Capacity cDNA Reverse Transcription Kit (Applied Biosystems, 4374966). Quantitative PCR (qPCR) was performed using TaqMan probes for *ST6GALNAC1* (Applied Biosystems Hs00300842_m1) with *GAPDH* (Applied Biosystems, Hs00266705_gl) as the endogenous control. Reactions were run in a Rotor-Gene 6000, and relative gene expression was calculated using the ΔΔCt method.

### 2.10 Dendritic Cell Differentiation and Treatment

Healthy volunteers, over 18 years of age were included in this study. The project was approved by the Human Research Ethics Committee of the Federal University of Sao Joao del Rei, Campus Centro Oeste Dona Lindu (CEPES CCO). All participants provided written informed consent prior to inclusion. Peripheral blood (30 mL) was collected to heparin tubes, and the peripheral blood mononuclear cells (PBMCs) were isolated by density gradient centrifugation using Ficoll-Paque Plus (GE Healthcare, USA), following the manufacturer’s instructions. The final cell pellet was collected; the cells were counted and resuspended in RPMI-1640 medium supplemented with 10% uFBS. Approximately 2×10^6^ cells/mL/well were plated in 24-well plates. The PBMCs were kept in serum-free RPMI□ 1640containing antibiotic□ antimycotic solution (100 U/mL penicillin, 25 μg/mL streptomycin, and 25 μg/mL amphotericin; Thermo Fisher Scientific Gibco, USA) at 37 °C and 5% CO2. Non-adherent cells were removed, and the adherent monocytes were harvested, washed,and cultured in fresh RPMI-1640 medium containing GM□ CSF (50 ng/ml; Immunotools, Germany), IL□ 4(50 ng/mL; Immunotools, Germany) and EVs (30 µg/mL) derived from MDA-MB-231-Mock or MDA-MB-231-STn culture supernatant for 5 days. DCs differentiated without EVs were used as controls. DCs were then activated with LPS (50 ng/mL; *Escherichia coli* 0111:B4; Sigma□ Aldrich, USA) for 48 h to induce maturation and subjected to phenotypical analysis or used in mixed lymphocyte reaction (MLR).

### 2.11 Dendritic Cells Phenotypic Analysis

DCs were differentiated from monocytes for 5 days and matured with LPS for 48h in the presence or absence of EVs. On day 7, DCs treated with EVs and untreated controls were analysed by flow cytometry to assess phenotypic changes. DCs were stained with the LIVE/DEAD™ Fixable Violet Dead Cell Stain (Invitrogen, USA) for 30 min according to the manufacturer’s instructions, and surface staining was performed using fluorochrome-conjugated antibodies against human HLA-DR (BioLegend, Cat No. 307610, 1:100), CD40 (BioLegend, Cat. No. 334310, 1:50), CD86 (BioLegend, Cat. No. 374204, 1:50) and CCR7 (BioLegend, Cat. No. 353216, 1:50). Flow cytometry acquisition was performed on an LSRFortessa™ X-20 flow cytometer with the software FACS Diva (BD Biosciences, USA). Data was analysed with Flow Jo 10.7 software (BD Biosciences, USA). The gating strategy involved selecting single cells, followed by gating on live cells (LIVE/DEAD-negative) and identifying DC. Expression levels of HLA-DR, CD40, CD86 and CCR7 were analysed within the live DC population.

### 2.12 Proliferation Assay

PBMCs obtained from healthy donors, as previously described, were used for isolation of allogeneic T lymphocytes. PBMCs were stained with anti-human CD3, CD4, and CD8 antibodies and sorted on a FACSAria II cell sorter (BD Biosciences, USA) to isolate highly purified CD4□ and CD8□ T-cell subsets (>95% purity). Each isolated allogeneic T lymphocyte subset was washed with PBS, counted, and labelled with CellTrace^TM^ carboxyfluorescein succinimidyl ester (CFSE, Invitrogen, USA) according to the manufacturer’s instructions. A MLR was set in the following way: DCs differentiated in the presence of different EVs were co-cultured with each isolated allogeneic T lymphocyte subset at a 1:10 DC:T-cell ratio in 96□ well U□ bottom plates and incubated at 37°C with 5% CO□ for 5 days. At the end of culture, cells were harvested and stained with LIVE/DEAD™ Fixable Violet (Invitrogen, USA, L34964) for viability assessment, followed by surface staining. Surface marker staining was performed by incubating the cells with monoclonal antibodies against human CD3 (Biolegend, 317324, V655, 1:40), CD4 (eBioscience, 48004842, V450, 1:160), CD8 (Biolegend, 301038, BV570, 1:20,000), CD25 (Biolegend, 302632, V605, 1:20), and CD127 (Biolegend, 351350, APC/Cy7, 1:50) for 30 minutes at 4°C in the dark in FACS buffer (PBS supplemented with 2% FBS and 2 mM EDTA). After staining, cells were washed twice with FACS buffer and resuspended for intracellular staining or flow cytometric acquisition.

For intracellular staining, cells were fixed and permeabilized using the FoxP3/Transcription Factor Staining Buffer Set (eBioscience™) according to the manufacturer’s instructions, and incubated with anti-human FoxP3 antibody (Biolegend, 320126, PE/Dazzle 594, 1:20) for 45 minutes at 4°C in the dark. Cell sorting was performed in LSRFortessa™ X-20 (BD Biosciences. Singlet lymphocytes were selected using the Boolean gating function (’Make and Gate’). Viable CD3□ lymphocytes (CD3□ ViViD□) were identified, followed by gating of CD4□ and CD8□ T lymphocyte cell subsets. Within CD4+, populations regulatory T cells were defined as CD25highdimCD127lowFoxP3□ For each T lymphocyte subset, proliferation was assessed based on CFSE dilution (CFSEdim).

### 2.13 Statistical Analysis

All experiments were performed using biological triplicates (n=3 independent donors for DC experiments; n=3 independent cell culture preparations for cell-based assays), with technical triplicates for each biological replicate. Data are expressed as mean ± standard deviation (SD). All statistical analysis were performed using GraphPad Prism version 9.0 for Windows (GraphPad Software, San Diego, California USA). Normality was assessed using Shapiro-Wilk test. For normally distributed data, statistical significance was determined using unpaired Student’s t-test (two groups) or one-way ANOVA with Tukey’s post hoc test for multiple comparisons. For non-normal data, Mann-Whitney U test or Kruskal-Wallis test was applied. A p-value of <0.05 was considered statistically significant.

## 3 Results

### 3.1 MDA-MB-231 Cells Overexpressing STn Produce a Higher Number and Larger Extracellular Vesicles (EVs)

In a recently published study in a cohort of 126 Triple Negative Breast Cancer (TNBC) patients, STn expression (in 23.8% of tumours) correlated to poorer survival (Supplementary Figure S1). To determine whether EVs from cancer cells expressing STn were also STn+, we used the TNBC MDA-MB-231 cell line transduced to overexpress *ST6GALNAC1* (MDA-STn). By flow cytometry, we confirmed that MDA-STn expressed the STn antigen at the cell surface (Figure 1A), and immunofluorescence imaging showed strong accumulation of STn near the nucleus and cytoplasmic staining, indicating its distribution in intracellular compartments (Figure 1B). These data, collectively, confirm the establishment of a robust STn+ tumour cell model for downstream EVs functional studies. To assess the quality, purity, and structural integrity of the isolated EVs, nanoparticle tracking analysis (NTA), and transmission electron microscopy (TEM) were performed. NTA revealed a monodisperse distribution of particles ranging from 30 to 150 nm across all samples, with a peak at around 100 nm, consistent with the physical characteristics of EVs. NTA photomicrographs visually confirmed the presence of vesicles in the expected size range **(Figure 1C-F).** TEM showed cup-shaped vesicles with double lipid bilayers, a clear indication of EVs, particularly given their endosomal origin. Negative staining with uranyl acetate clearly delineated round vesicles measuring 30-120 nm in diameter, corroborating the structural identity of purified EVs (Figure 1F).

**Figure 1.**
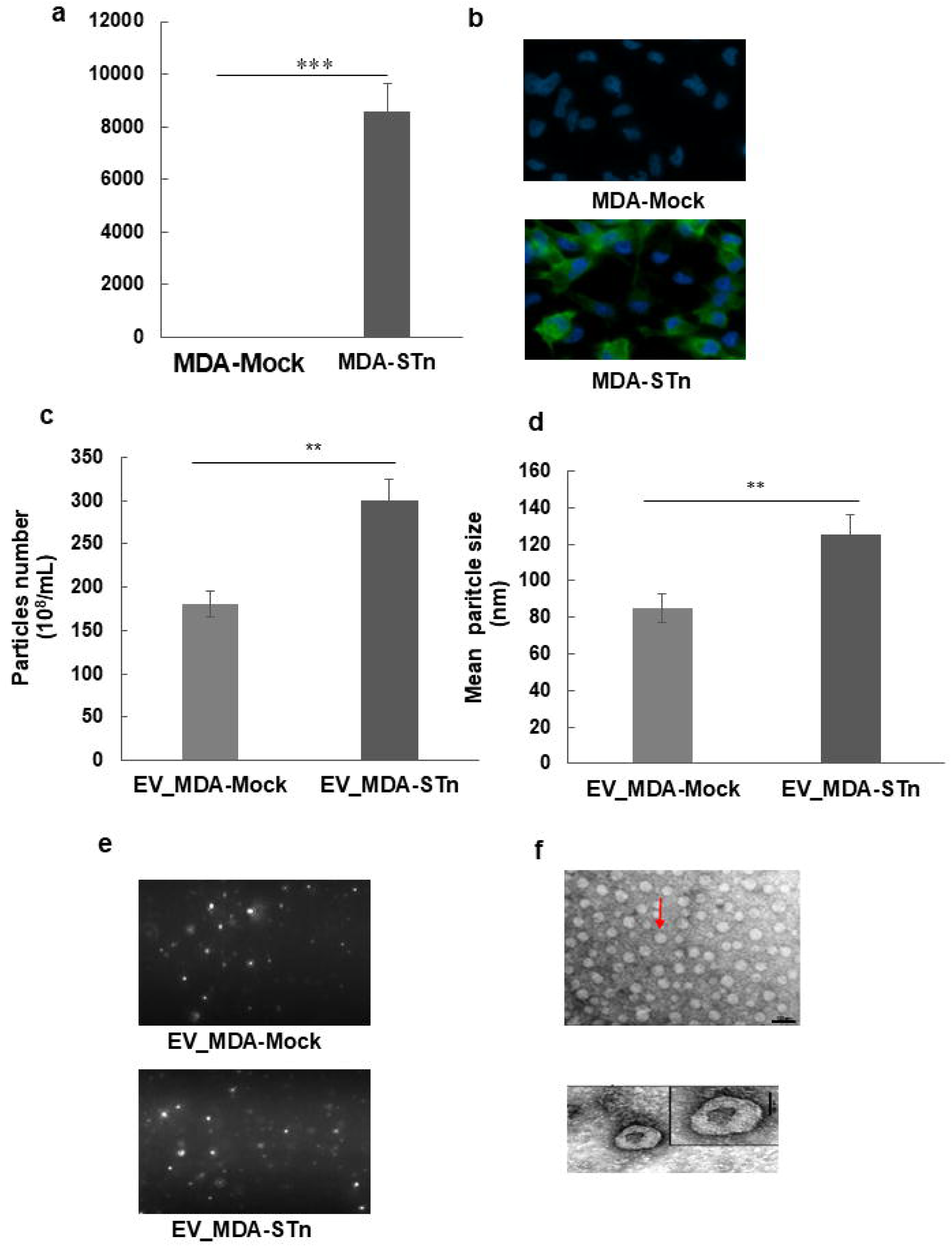
Characterization of EVs derived from MDA-MB-231 overexpressing STn. STn expression was assessed in MDA-MB-231 Mock and STn□ cells by (a) flow cytometry and (b) Immunofluorescence microscopy demonstrating the presence of STn in cells transduced with ST6GalNAc-I.Nanoparticle tracking analysis (NTA) performed on cell culture supernatants showed that (c) STn□ cells released a significantly higher number of particles with (d) increased average diameter compared to Mock cells. (e) Images of nanoparticle tracking analysis (NTA). (f)Transmission electron microscopy revealed the presence of small vesicles with characteristic cup-shaped morphology in both groups. Data represents at least three independent experiments and are expressed as mean ± SEM. Statistical significance was determined using unpaired Student’s t-test: \**p* < 0.05, \*\**p* < 0.01, \*\*\**p* < 0.001.

Regarding the concentration and mean particle size of the isolated EVs, the data revealed a significantly elevated particle concentration for EV released from MDA-STn, with a peak particle concentration of approximately 300 ± 9.5×10□ particles/mL compared to 180 ±8.3× 10□ particles/mL from the Mock control (Figure 1C). In parallel, mean particle size differed significantly between the two conditions. EVs from MDA-MB-231-STn cells exhibited a mean diameter of 125 ±7.4 nm with uniform distribution compared to a mean diameter of 85 ±9.8 nm for the Mock control (Figure 1D). These results we corroborated with the microscopic images taken during NTA (Figure E,F). These morphological observations corroborate the quantitative results, indicating the alterations in EV production and structure associated with the aberrant STn glycan expression

### 3.2 Extracellular Vesicles from STn+ MDA-MB231 Cell Line are Rich in STn and also Incorporate ST6GalNAc1 mRNA

Immunoblot analysis of lysates from EVs and donor cells revealed the presence of STn antigen in vesicles derived from MDA-STn cells. Sialidase treatment efficiently removed terminal sialic acid residues, eliminating STn signal, confirming the glycan’s identity. STn expression was undetectable in mock-transduced controls, also corroborating the identity of the glycan (Figure 2A, left panel). STn-positive extracellular vesicles (STn+ EV) were detected in supernatants collected after 48h of cell culture, with consistent enrichment of CD9 and CD81, indicating that EVs were reliably isolated during the process (Figure 2A, right panel). These findings indicate that tumour cells can package STn-carrying glycoproteins into EVs, suggesting a functional delivery mechanism of tumour-associated glycoantigens. To assess the specificity and efficacy of glycoengineering in MDA-MB-231 cells and evaluate glycosyltransferase transcript incorporation, qRT-PCR was performed to quantify ST6GalNAcI expression in mock- and ST6GalNAcI-transduced (MDA-STn) cells. As expected, ST6GalNAc-I expression was significantly upregulated in MDA-STn cells compared to Mock controls (*p*< 0.0001), confirming the efficacy of lentiviral transduction (Figure 2B). The ST6GalNAcI mRNA incorporation into EV of MDA-STn was also assessed by qRT-PCR. ST6GalNAc-I was also detected in EVs derived from STn□ cells, although at lower levels than in the corresponding producer cells (Figure 2B), suggesting that a fraction of the transcript was selectively incorporated into EVs. This observation may indicate a potential route for intercellular transfer of glycosylation-related transcripts within the tumour microenvironment, although further studies are required to confirm this mechanism.

**Figure 2.**
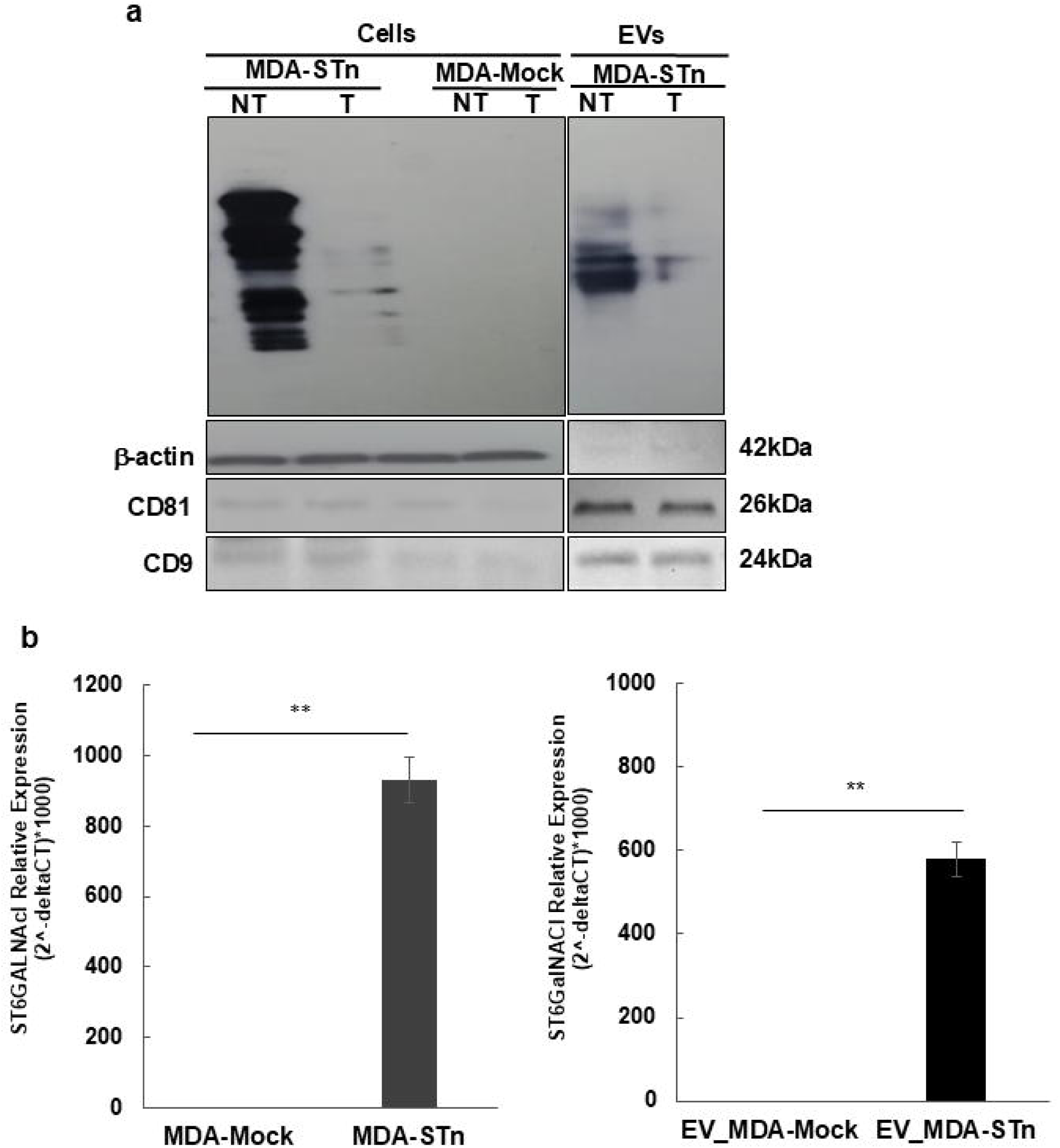
Detection of STn and ST6GalNAc-I expression in EVs derived from STn+ MDA-MB-231 cells. (a) Western blot analysis to detect STn expression in total protein lysates from MDA-MB-231-STn (MDA-STn), MDA-Mock, and in EVs isolated from MDA-STn cells. Lysates were either not treated (NT) or treated with sialidase (T). β-actin was used as loading control for cell lysates, CD81 and CD9 were used as EV markers and loading controls. (b) Quantitative PCR analysis of ST6GalNAc-I mRNA expression in MDA-MB-231-STn and Mock-transfected cells (left), and in their respective EVs (right). Results are shown as relative expression (2□ ΔCt × 1000), normalized to GAPDH. Data represent mean ± SD from three independent experiments. Statistical significance was determined using unpaired Student’s t-test: \*\**p* < 0.01.

### 3.3 STn□ EVs Mediate the Transfer of STn Antigen to DCs and Impaired Maturation

To study the effect of EVs, monocytes were cultured in differentiation medium containing EVs isolated from MDA-MB-231 Mock transduced cells (EV_MDA-Mock), STn□ cells (EV_MDA-STn), STn□ EVs pre-treated with sialidase (EV_MDA-sialidase). On day 5, immature dendritic cells (iDCs) were challenged with LPS to induce their maturation and compare the effect on DCs incubated with LPS alone (mDCs) with those exposed to EVs with differential glycosylation, on the expression of the maturation markers, CD86, HLA-DR, CCR7, and CD40. (Figure 3A). In Figure 3B we show the gating strategy for flow cytometry analysis: singlets were selected based on SSC-H/SSC-A, followed by identification of the DC population by granularity and size, and gating of live cells using ViViD dye exclusion. DĆs viability after EV treatment remained high across all experimental conditions (>93% viable cells) (Figure 3C), indicating that subsequent phenotypic differences were not attributable to cytotoxicity DCs cultured in the presence of EV_MDA-Mock from already displayed an impaired maturation profile, with significant lower upregulation of all the above-mentioned markers than DCs. . Interestingly, DCs cultured with EV_MDA-STn exhibited complete maturation abrogation, with near-complete absence of CCR7 and HLA-DR expression and significantly decreased levels of CD40 and CD86 compared to DCs cultured with EV_MDA-Mock and to mDCs. Strikingly, this suppressive phenotype was reversed upon sialidase treatment of STn□ EVs prior to DC exposure. DCs incubated with EV_MDA-sialidase exhibited a maturation profile comparable to or greater than that of Mock-EV-treated cells, with partially restored MFI levels for CD86, HLA-DR, CCR7, and CD40, highlighting the role played by the sialylation status of EVs—specifically the STn epitope—in suppressing DCs maturation.

**Figure 3.**
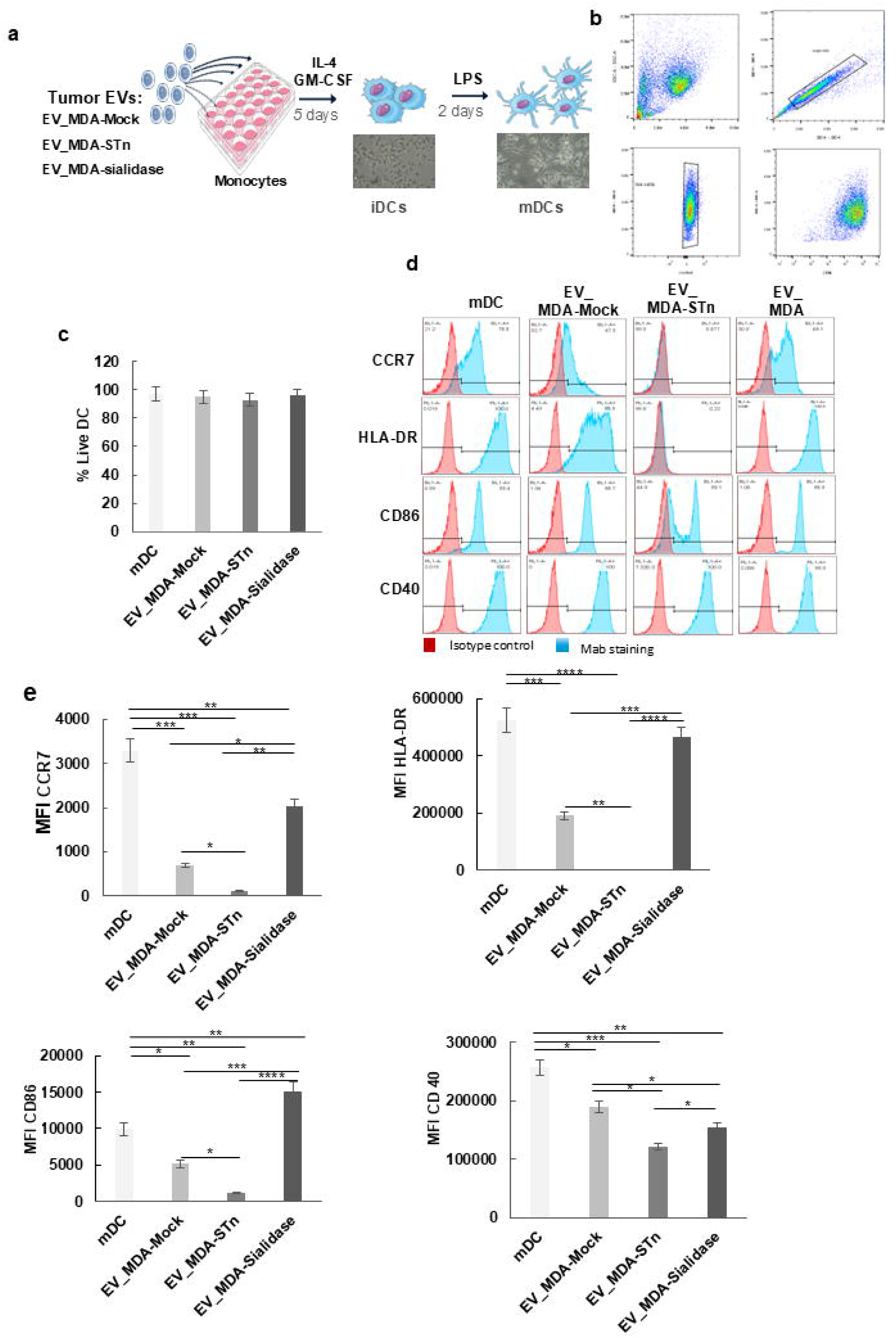
Expression of maturation markers in DCs was reduced when cultured in the presence STn+-EVs. (a) Schematic of experimental workflow for comparing the effect of different EVs on DC maturation. DCs were generated from monocytes by culturing with IL-4 and GM-CSF for 5 days in the absence or presence of EVs obtained from MDA-MB-231 cells: mock-transduced (EV_MDA-Mock), transduced to overexpress STn (EV_MDA-STn), or STn□ EVs treated with sialidase (EV_MDA-STn-Sialidase). On day 5, cells were stimulated with LPS for 48 h to induce maturation. (b) Gating strategy for flow cytometry analysis: singlets were selected based on SSC-H/SSC-A, followed by identification of the DC population by granularity and size, and gating of live cells using ViViD dye exclusion. (c) Evaluation of DC’s viability(d) Representative histograms of surface expression of the maturation markers CCR7, HLA-DR (MHC-II), CD86 and CD40 in DCs. Red histograms represent isotype controls and blue histograms indicate specific staining (e) Quantification of the median fluorescence intensity (MFI) of expression of maturation markers. Results represent three independent experiments (mean ± SD). Statistical analyses were performed using one-way ANOVA followed by Tukey’s post hoc test: \**p* < 0.05; \*\**p* < 0.01; \*\*\**p* < 0.001; \*\*\**p* < 0.0001.

Collectively, these findings indicate that EVs carrying STn glycoforms from MDA-MB-231 cells can inhibit the phenotypic maturation of monocyte-derived dendritic cells in vitro. The reversibility of this effect through enzymatic desialylation underscores the glycan-dependency of this immunosuppressive mechanism and highlights glycoengineering as a key modulator of immune cell function in the tumour microenvironment.

### 3.4 STn+ EV-exposed DCs Caused Impairment of T cell Proliferation

To investigate the functional consequences of DC conditioning by tumour-derived EVs, we performed a mixed lymphocyte reaction (MLR) assay with DCs generated in the presence of EVs isolated from MDA-MB-231 mock-transduced and MDA-MB-231-STn, as well as STn□ EVs treated with sialidase. The ability of EV-primed DCs to influence the proliferation of allogeneic CD4□, CD8□, and regulatory (Treg) T cells was evaluated by CFSE dilution, as detected by flow cytometry (Figure 4A).

**Figure 4.**
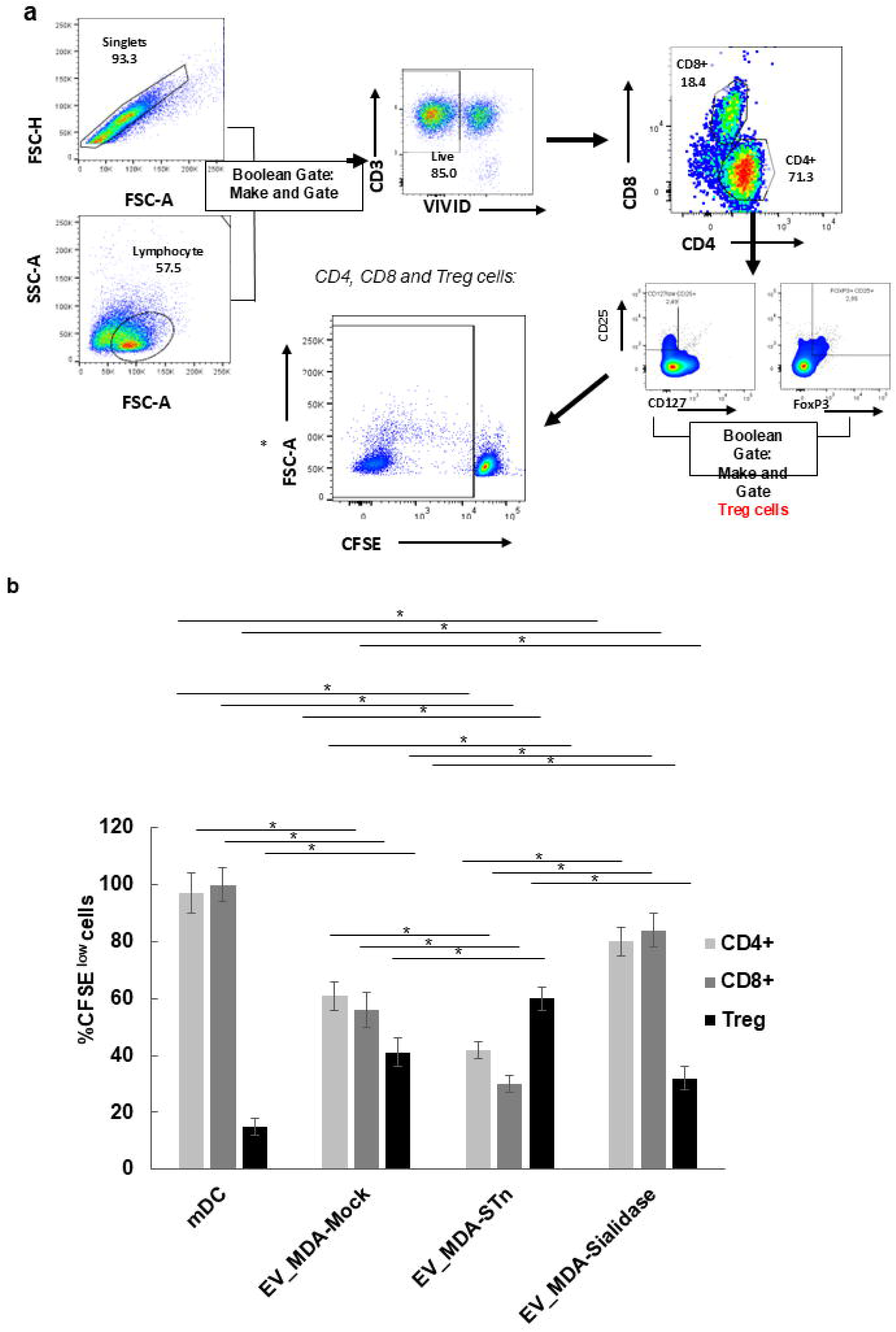
STn+ EVs reduce DĆs function and T cell proliferation. (a) Gating strategy used for flow cytometric cell sorting and proliferation analysis of CD4□, CD8□, and regulatory T cells (Tregs) Singlets were identified by FSC-A vs FSC-H, and the lymphocyte population was gated based on FSC-A vs SSC-A. Viable T cells were selected as CD3□ ViViD□ CD4□ and CD8□ subsets were then sorted. Regulatory T cells were identified within the CD4□ gate using a Boolean combination of CD25^high^CD127^low^ and CD25□ FoxP3□ expression. After a mixed lymphocyte reaction (MLR) co-culture for 5 days, proliferation was determined by CFSE dilution (CFSE^dim^) (b) Proliferation of sorted allogeneic CD4□, CD8□, and Treg subsets co-cultured with monocyte-derived dendritic cells (DCs) LPS-treated DCs never exposed to EVs (mDC) or DCs in the presence of EVs from MDA-MB-231 Mock (EV_MDA-Mock), MDA-MB-231 STn□ ((EV_MDA-STn), or EVs from MDA-MB-231 STn□ cells treated with sialidase (EV_MDA-STn-Sialidase). Data represent mean ± SD from three independent experiments. Statistical significance was determined using one-way ANOVA followed by Tukey’s post hoc test: \**p* < 0.05.

EVs from MDA-Mock-transduced cells caused a significant 36 ± 4% reduction in proliferative CD4□ T cells and 44 ± 6% reduction in proliferative CD8□ T cells, together with 26 ± 5% increase in proliferative CD4□ CD25□ FoxP3□ Treg cells. These effects were more pronounced for STn EVs-conditioned DCs, which suppressed 55 ± 6% of CD4□ T cell proliferation, 70 ± 6% of CD8□ T cells proliferation and increased 45 ± 4% Treg cell expansion. Compared to the MOCK-treated condition, the reduction in proliferation was 19% greater for CD4□ T cells, 26% greater for CD8□ T cells, while Treg expansion increased 19%. These effects were abrogated by sialidase treatment of STn□ EVs, restoring DC-driven T cell activation and reducing Treg induction. Combinedly, these results support that the immunomodulatory properties of tumour EVs depend on their glycan composition. Specifically, terminal sialylation of STn□ EVs appears critical for promoting tolerogenic DC–T cell interactions.

### 3.5 EV *cargo* Mediated the Transfer of STn Antigen and STn-Synthesizing Enzyme to DCs

To assess the ability of DCs to incorporate STn antigen and ST6GalNAc-I enzyme from EV’s cargo, they were incubated under four conditions: DCs alone, DCs + EV_MDA-Mock, DCs + EV_MDA-STn, and DCs + EV_MDA-STn pre-treated with sialidase for 48h.Western blot analysis revealed a strong ST6GalNAc-I signal in lysates from MDA STn cells and their derived EVs (Fig. 5A, lanes 1 and 2, respectively), showing successful overexpression and active sorting of the enzyme into the vesicular cargo. Strikingly, ST6GalNAc-I was also detected in DCs exposed to STn□ EVs (Fig 5A, lane 3). In contrast, the enzyme was absent in DCs exposed to vesicles from MDA-Mock (Fig 5A, lane 4) or untreated DCs (Fig 5A, lane 5), and sialidase treatment had no influence on the transfer of ST6GalNAc-I Fig 5A, lane 6), providing compelling evidence that tumour EVs can deliver functional glycosylation enzymes to dendritic cells. To determine whether glycan transfer accompanied enzyme delivery, flow cytometry was used to quantify STn antigen acquisition on DCs’ surfaces. While untreated and mock-EV–treated DCs showed negligible STn signal, a significant increase in STn median fluorescence intensity (MFI) was observed in DCs incubated with STn□ EVs (Figure 5 B, C). Notably, this effect was abrogated when donor EVs were pre-treated with *Clostridium perfringens* sialidase, confirming that the transferred glycan was STn. These findings validate that STn□ EVs mediate the transfer of both the glycosyltransferase ST6GalNAc-I and its glycan product.

**Figure 5.**
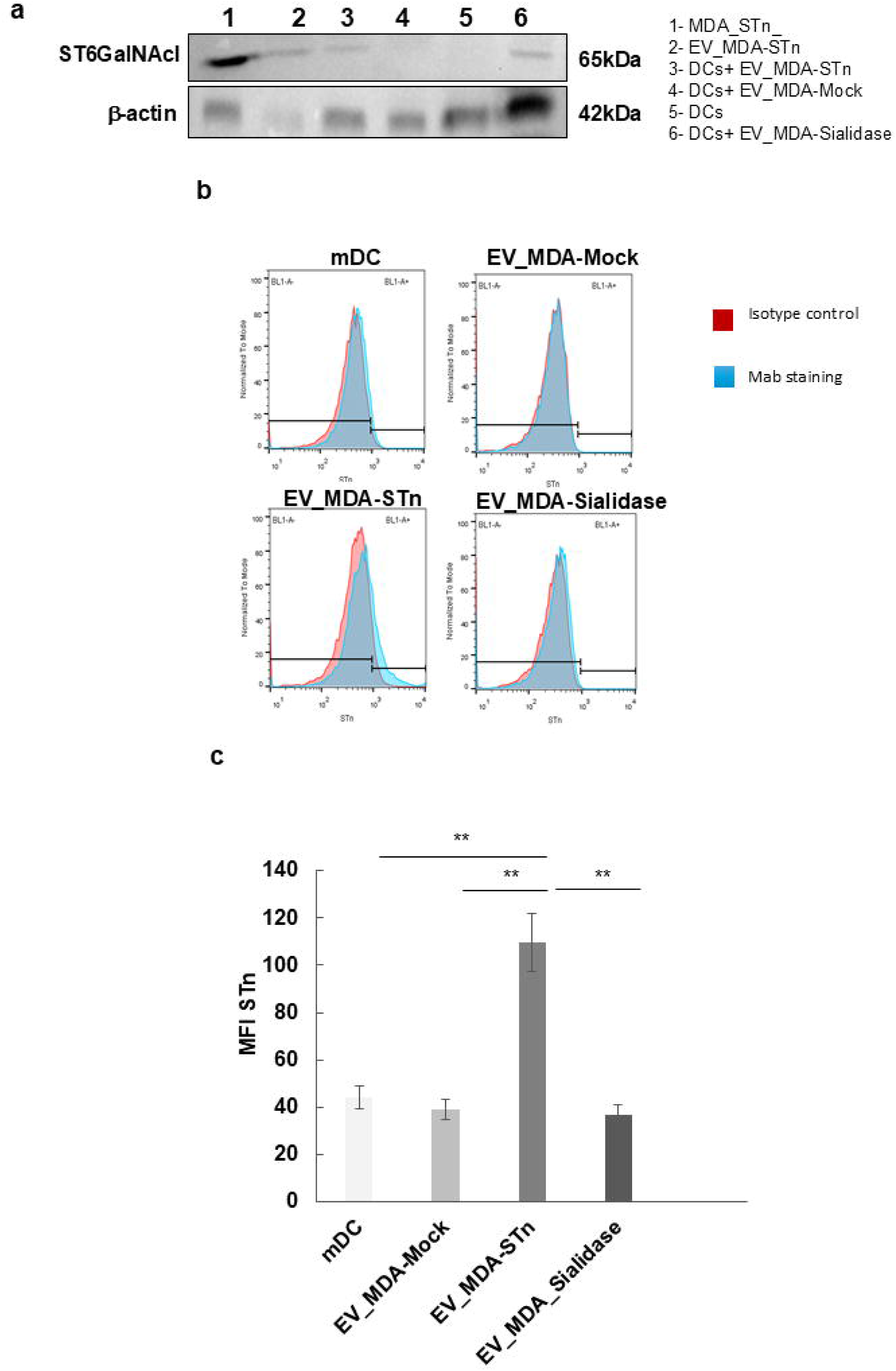
STn+ EV are responsible for the transfer of both STn glycan and the STn synthesizing enzyme to DCs. (a) Immunoblot detection of ST6GalNAc-I in (1) lysate of MDA_STn_cells, (2) lysate of EV_MDA-STn, (3) lysate of DC treated with EV_MDA-STn, (4) lysate of DC treated with EV_MDA-Mock (5) lysate of DC without treatment and (6) lysate of DC treated with EV_MDA-Sialidase. (b) Representative histograms of STn expression in DCs incubated under four conditions: DCs alone (top left), DCs + EV_MDA-Mock (top right), DCs + EV_MDA-STn (bottom left), and DCs + EV_MDA-STn pre-treated with sialidase (bottom right). Isotype control (red line) and anti-STn staining (blue filled histogram) are shown. (c) Quantification of the median fluorescence intensity (MFI) of STn expression. Data represent mean ± SD from three independent experiments. Statistical significance was determined using, one-way ANOVA with Tukey’s multiple comparison test: \*\**p* < 0.01

Together, these findings reveal that tumour-derived EVs not only carry bioactive proteins and RNA but can function as vehicles of glycosylation enzymes capable of reshaping immune cell phenotypes.

## 4 Discussion

The immunological crosstalk between tumours and dendritic cells (DCs) constitutes a dynamic and clinically relevant area of cancer immunology. Tumour-derived extracellular vesicles (TDEVs) are increasingly recognised as immunoregulatory mediators capable of altering the functional behaviour of recipient immune cells. For instance, studies have reported that TDEVs can influence DC antigen uptake and intracellular processing (52). Aberrant glycosylation is a prominent feature of cancer cells that spreads to EVs produced by these cells (53,54), with cancer-associated STn glycoepitope is often found decorating tumour proteins (55). However, the contribution of defined glycan structures to the immunomodulatory functions TDEVs remains poorly characterised. In this study, we demonstrate that EVs enriched in the cancer-associated STn glycoepitope modulate DC immunogenicity, promoting a tolerogenic phenotype shift that may facilitate tumour immune evasion. This phenotype is characterised by reduced expression of CD86, CD40, CCR7 and HLA-DR, impaired T-cell priming capacity, and enhanced expansion of regulatory T cells. Notably, these effects occurred in the absence of detectable cytotoxicity, indicating that DC modulation is mediated by molecular cues, specifically tumour-associated glycans, rather than by EV-induced cell damage. Importantly, our model employs MDA-MB-231 cells stably transduced with the glycosyltransferase ST6GalNAc-I, which catalyzes the biosynthesis of STn. This system is biologically and clinically relevant. Clinically, STn is aberrantly expressed in a significant subset of primary breast carcinomas. In a cohort of patients with invasive breast cancer, STn was detected in approximately 39% of tumours and was associated with estrogen/progesterone receptor negativity and reduced 5-year survival (56). Subsequent reviews confirm that 20–30% of breast tumours express STn, with variability depending on detection method and tumour subtype (57). Furthermore, experimental studies confirm that enforced ST6GalNAc-I expression in breast cancer cells induces STn surface presentation and increases tumourigenicity and invasiveness in vivo (58). Collectively, these findings support the translational relevance of our glycoengineered EV model in recapitulating tumour-specific glycosylation patterns observed in clinical diseases (59). Tumor-derived EVs have been shown to induce tolerance and, thus, promoting tumor growth and metastasis by different mechanisms, one of them being imposing an immunosuppressive environment (60). Not surprisingly, the EVs derived from MDA-MOCK already cause a significant impairment of dendritic cell maturation, most likely driven by tumor-derived immunosuppressive signals carried by these vesicles, encompassing lipids, proteins, nucleic acids, and metabolites, as reviewed (61). But, strikingly, in this work, we show that the presence of STn significantly reinforces the immunosuppressive effect, evidencing its specific role in the suppression mechanism. The binding of STn decorated proteins to Siglec receptors on the surface of DCs is the most likely event that account for STn dependent immunosupression caused by STn +EVs as shown for (62,63)

In addition to altering DC maturation and T□ cell priming, STn□ EVs promoted the appearance of STn on recipient DCs, consistent with horizontal transfer of tumour□ associated glycan features. This observation aligns with emerging evidence that EV cargo can include glycosylation machinery capable of modifying recipient cells after uptake. In our system, STn□ EVs transferred ST6GalNAc□ I to DCs and were associated with STn acquisition on the DC surface, supporting a model in which tumour EVs can disseminate glycan determinants and glycosyltransferases within the tumour microenvironment. This mechanism provides a potential explanation for how tumour□ associated sialoglycans reinforce tolerogenic DC programming and contribute to immune escape.

Further reports show that exosomal glycosyltransferases can be enzymatically active upon uptake by target cells. Liang et al. demonstrated that exosomal FUT8 transferred from hepatocellular carcinoma cells to fibroblasts catalyzed core fucosylation in recipient cells, promoting a tumour-supportive phenotype (64). Similarly, Kore et al. reported that β1,6-N-acetylglucosaminyltransferase V (GnT-V) delivered via melanoma EVs enhanced β1,6-branching in endothelial cells and promoted angiogenesis (65). Also in aggressive prostate cancer, recent evidence points to ST6GAL1transfer by small EVs to potentially modulate cell surface sialylation in recipient cells and promote bone marrow tropism and metastatic progression (66) These studies validate the concept that EVs can deliver functionally competent glycosylation enzymes rather than merely static glycan residues to modulate host cell identity and function.

Consistent with these findings, we detected ST6GalNAc-I RNA transcripts in recipient DCs, evidencing the horizontal transfer of glycosylation machinery itself. Moreover, the levels of related sialyltransferase ST6GalNAc-II RNA transcript, which was endogenously expressed in mock-transduced MDA-MB-231 cells suffered a slight reduction in MDA-MB-231-STn, suggesting regulatory adjustments following ST6GalNAc-I overexpression. A similar pattern for these ST6GalNAc RNA transcripts was observed for EV derived from the respective cells (results not shown). This suggests that exosome-mediated dissemination of enzymatic instructions may underlie immune cell reprogramming, a mechanism warranting deeper investigation in glycoimmunology.

Importantly, the immunosuppressive activity of STn□ EVs was fully reversed by treatment with *Clostridium perfringens* sialidase, indicating that terminal sialic acid residues are essential for suppression of DĆs function. These data align with prior reports demonstrating that sialylated glycans engage inhibitory Siglec receptors on DCs, blunting their maturation and antigen-presenting function (67,68). Indeed, we recently demonstrated that sialic acid alone is sufficient to drive tolerogenic DC programming (69), reinforcing the central immunomodulatory role of this glycan class. In line with this, enzymatic removal of sialic acid from dendritic cells was shown to enhance cross-presentation capacity, upregulate co-stimulatory molecules (CD86, CD40), and promote robust CD8□ T cell activation and IFN-γ secretion, as previous demonstrated (69,70). These findings support a model in which tumour-associated sialoglycans, such as STn, promote immune tolerance by engaging inhibitory siglec receptors—mechanisms that can be reversed through targeted glycan editing (40,69–71).

Our findings have important therapeutic implications. First, the demonstration that sialidase treatment reverses DC suppression suggests that targeting tumour-associated sialoglycans may restore anti-tumour immunity. Second, STn□ EV levels in patient serum could serve as a liquid biopsy biomarker for immune evasion status. Third, blocking EV uptake or glycosyltransferase transfer may represent novel immunotherapeutic strategies in STn breast cancers. However, several limitations warrant consideration. First, our in vitro model using MDA-MB-231 cells may not fully recapitulate the complexity of the tumour microenvironment. Second, while we demonstrate glycan and enzyme transfer, the precise molecular mechanisms of EV uptake and intracellular trafficking remain to be investigated. Third, validation in primary human breast cancer samples and in vivo models is needed to confirm clinical relevance. Finally, the relative contributions of direct glycan recognition versus de novo glycosylation in recipient cells remain to be fully elucidated.

Together, our findings support a model in which tumour-derived EVs enriched in STn glycoepitopes suppress DC immunogenicity through dual mechanisms: direct engagement of Siglec receptors via sialylated glycans, and the delivery of glycosylation-modifying cargo that may alter recipient cell glycosylation (Fig. 6). These insights reinforce the concept that the tumour glycome extends beyond the cancer cell surface to shape the systemic immune landscape via extracellular vesicles (72,73).

**Figure 6.** Schematic representation of the STn glycan horizontal transfer to dendritic cells and of its negative impact DĆs function and T cell priming. Tumour-derived EVs enriched in STn glycoepitopes suppress DC immunogenicity through dual mechanisms: direct engagement of Siglec receptors via sialylated glycans, and the delivery of glycosylation-modifying cargo that may alter recipient cell glycosylation (image was generated in BioRender: Scientific Image and Illustration Software, https://www.biorender.com/).

By leveraging a glycoengineered system with high translational relevance, we provide mechanistic insight into how glycosylated EVs contribute to immune escape. These results not only extend previous studies on glycan-mediated immune suppression (40,73,74) but also reveal potential therapeutic targets, including glycan editing and inhibition of EV communication pathways.

## Supporting information

graphical abstract

## Author Contributions

N.D.A. contributed to experiments performance, analysed data, conceptualization, interpretation, figure construction, coordination, manuscript writing, reviewing and funding acquisition. S.S. contributed to data analysis, interpretation, figure construction, review and manuscript writing. M.S.V. and B.R.O. contributed to experiment performance. Z.S. contributed to the study rationale, data analysis, interpretation, figure construction and manuscript writing. P.A.V. contributed to conceptualization, coordination and funding acquisition. All authors read, revised and approved the final manuscript.

## Acknowledgements

The authors would like to thank the FEI Company at the Electron Microscopy Center of UNESP/Botucatu and Egas Moniz University, Caparica, Portugal, for providing access to the transmission electron microscopy facilities and for their valuable technical support. We are also grateful to the technicians Elisabete Ferreira and Cláudia Carvalho for their kind assistance with centrifugation and ultracentrifugation procedures. **Funding:** This work was supported by the FAPEMIG (Grant APQ-02663-16). Additionally, a Master’s scholarship was provided to Brenda Raissa de Oliveira from CAPES. This work was supported and under grants UIDP/04378/2020 and UIDB/04378/2020 (provided to the Applied Molecular Biosciences Unit – UCIBIO), LA/P/0140/2020 (provided to the Associate Laboratory Institute for Health and Bioeconomy – i4HB). by Portuguese Foundation for Science and Technology (FCT) with project InnoGlyco (ref: 2022.04607.PTDC) This work was also supported by the European Commission through the GLYCOTwinning project (Grant Agreement: 101079417), and by the European Union’s Horizon 2020 research and innovation programme under the EJPRD COFUND-EJP N 825575 (EJPRD/0001/2020).

## Data Availability Statement

The data that support the findings of this study are available from the corresponding author upon reasonable request.

## Abbreviations

APCs: antigen-presenting cells
BCA: bicinchoninic acid
DCs: monocyte derived dendritic cell
DMEM: Dulbecco’s Modified Eagle’s Medium
ECL: enhanced chemiluminescence
EV: extracellular vesicles
FBS: fetal bovine serum
GnT-V: β1,6-N-acetylglucosaminyltransferase V
HEK: human embryonic kidney
MFI: median fluorescence intensity
MLR: mixed lymphocyte reaction
NTA: nanoparticle tracking analysis
PBSCs: peripheral blood mononuclear cells
PBS: phosphate buffered saline
RIPA: radioimmunoprecipitation assay buffer
SD: standard deviation
ST6GalNAc-I: N-acetylgalactosaminide alpha-2,6-sialyltransferase I
TCGA: The Cancer Genome Atlas
TDEVs: tumour-derived extracellular vesicles
TEM: transmission electron microscopy
TGF-β: Transforming growth factor-beta
TNBC: triple negative breast cancer
Tregs: regulatory T cells.

