## Supplementary material for "Sialyl-Tn-positive tumour-derived extracellular vesicles impair dendritic cell function via horizontal transfer of glycans": Fig S1

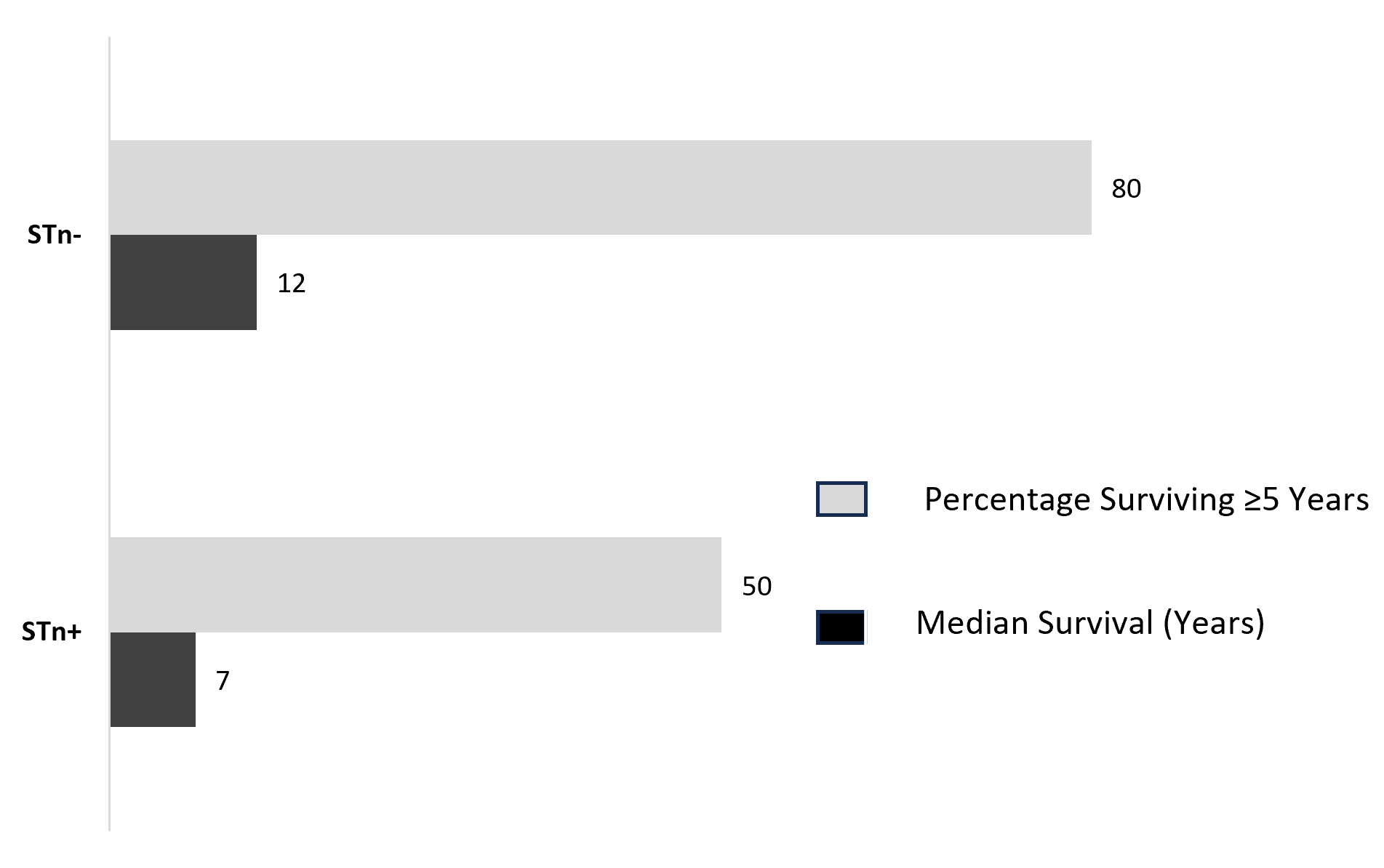

**Fig S1- STn expression is associated with the lower overall survival in the TNBC patients.** To characterise STn expression in TNBC cancer tissues tumour tissue microarrays (TMA) was performed. In a retrospective study of a cohort of 126 TNBC, the Kaplan-Meier method was used to analyse the overall survival in stratified STn+ and STn- groups. From the group of STn-patients, 80% survived at least 5 years, with a median survival time of 12 years, From the group of STn+ patients, 50% survived at least 5 years, with a median survival time of 7 years.
