## Supplementary figures and images for "Sialyl-Tn-positive tumour-derived extracellular vesicles impair dendritic cell function via horizontal transfer of glycans"

### graphical abstract

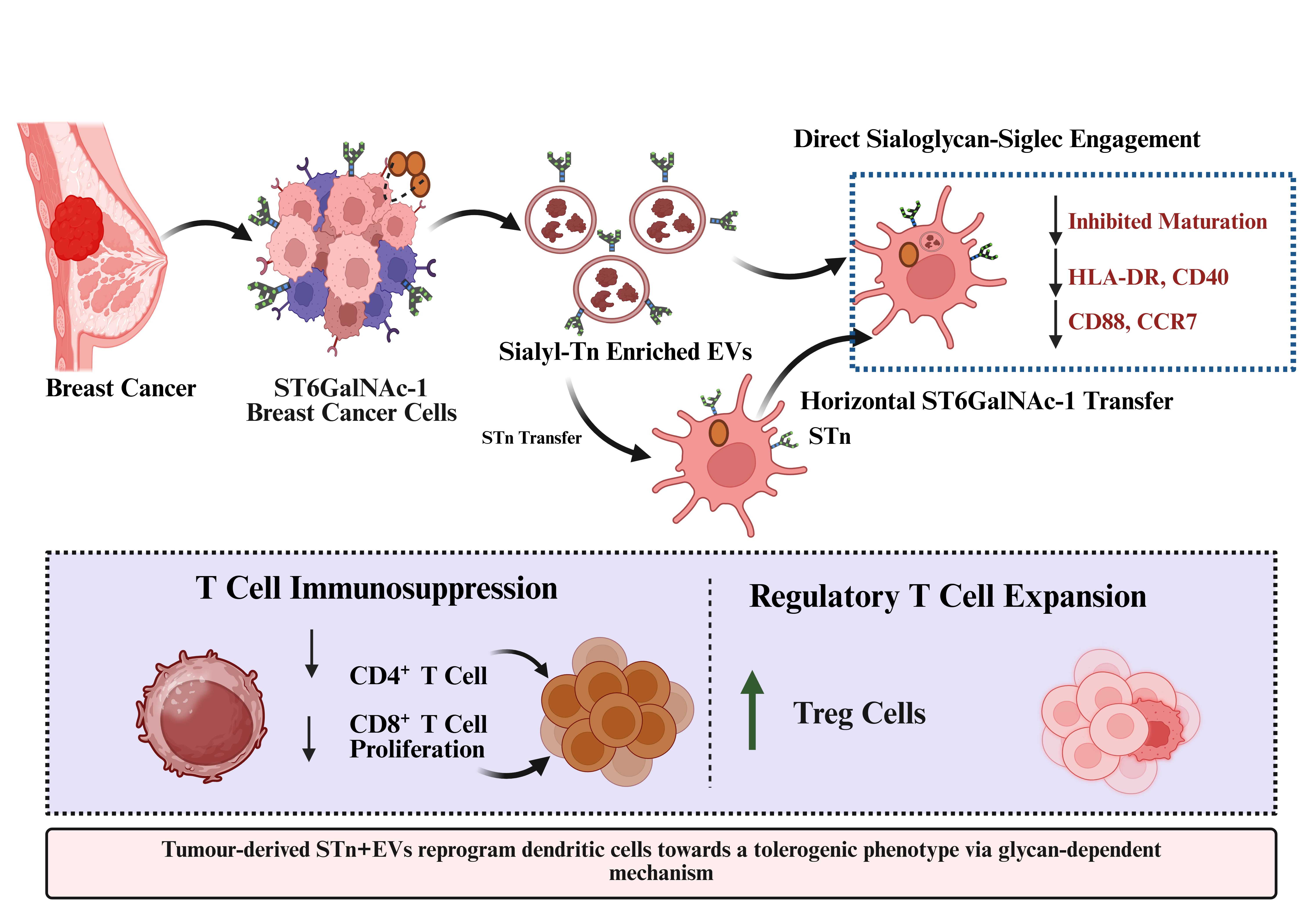
